# BioAct: Biomedical Knowledge Base Construction using Active Learning

**DOI:** 10.1101/2022.04.14.488416

**Authors:** Dustin Wright, Anna Lisa Gentile, Noel Faux, Kristen L. Beck

**Affiliations:** University of Copenhagen; IBM Research

## Abstract

Creating and curating knowledge resources has been a paramount activity in the biomedical domain. In recent years, automated methods for knowledge base construction have flourished and have enabled large scale construction and curation of such resources. In the biological domain, techniques such as *next generation sequencing* produce new data at exponential rate, making mere manual curation of knowledge resources simply unfeasible. The major technology to automate knowledge base construction is Information Extraction — specifically tasks such as Named Entity Recognition or Relation Extraction. The major hurdle for IE methods is the availability of labelled data for training, which can be prohibitively expensive and challenging to obtain due to the need of domain experts. Active learning aims at minimizing the cost of manual labelling by only requiring it for smaller and more useful portions of the data. With this motivation, we devised a method to quickly construct highly curated datasets to enable biomedical knowledge base construction. The method, named *BioAct*, is based on a partnership between automatic annotation methods (leveraging SciBERT with other machine learning models) and subject matter experts and uses active learning to create training datasets in the biological domain. The main contribution of this work is twofold; in addition to the *BioAct* method itself, we publicly release an annotated dataset on antimicrobial resistance, produced by a team of subject matter experts using *BioAct*. Additionally, we simulate a knowledge base construction task using the MegaRes and CARD knowledge bases to provide insight and lessons learned about the usefulness of the annotated dataset for this task.

## 1 Introduction

Training a deep learning model for automatic knowledge base construction requires access to massive stores of labelled data. However, in most scientific domains such data does not exist and would be expensive to obtain. With biomedicine in particular, the cost can be prohibitively expensive as labelling requires a high level of expertise [25, 18]. This prevents the adoption of certain tools, such as those for information extraction, in scientific fields which could benefit from them due to the massive amount of literature being published every day [38]. Given this, the primary question we are asking in this work is: how can one quickly create a tool for automatic biomedical knowledge base construction in a particular scientific domain starting with limited data?

To this end we introduce *BioAct*, a method for constructing biomedical knowledge bases starting from limited data, along with a case study dataset focusing on antimicrobial resistance generated using this method. *BioAct* relies on active learning to quickly construct the dataset while maintaining high quality data. We tie collecting data and training a machine learning model for both entity recognition and relation extraction into a single active learning pipeline. Unlabelled data for relation extraction is generated progressively from corrected entity predictions. We demonstrate that the labels created using this method continuously improve the ability of a model to augment an existing seed knowledge base through many iterations of active learning.

We focus on the particular domain of antimicrobial resistance [13], which has seen an explosion of literature in recent years as it has gained recognition as a global public health threat [30]. An example sentence containing information of interest in this domain is given in Figure 1. In order to provide better treatment plans or discover new effective drugs, antimicrobial resistance is of great interest to the biomedical community and having resources to quickly sift and search through the expanding literature will be a boon to researchers in the field. To date, there are a handful of manually curated knowledge bases containing facts related to antimicrobial resistance including CARD [16] and MegaRes [21]. Yet, the efforts to produce these are laborious and do not scale in-line with the rate of publications created describing bacterial genome evolution conferring new resistance mechanisms. Given this, one of our goals was to provide the resources necessary to automatically build a knowledge base of antimicrobial resistance facts and alleviate the need for manual curation.

**Figure 1.**
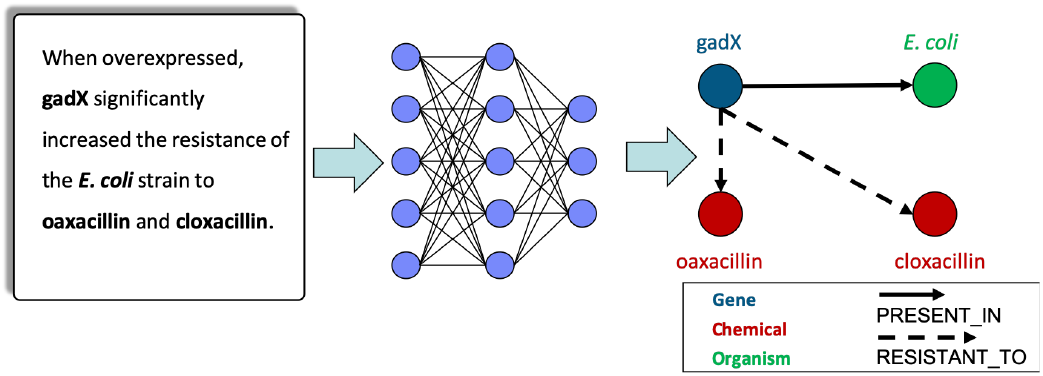
Automatic construction of a knowledge base of antimicrobial resistance facts

To evaluate the efficacy of *BioAct*, we performed a knowledge base completion task where we held out a portion of a known knowledge base with associated relevant documents to compare against. We show that a model trained on the labels acquired during active learning continuously improves in its ability to recover the knowledge base, as well as to augment it with new relations. In addition, we show that we can acquire useful training data using active learning with less annotation budget within this domain. We plan to release the dataset that we collected for the benefit of the BioNLP community.

The core contributions of our work are as follows:

- A method for quickly bootstrapping a tool for automatic knowledge base construction yielding twice as many labels with 25% of the amount of annotators over similar periods of time.
- A dataset labelled with entities and relations for extracting antimicrobial resistance facts.

Our experimental study provides insight into how a model trained for information extraction using active learning improves for the task of automatic knowledge base construction over several iterations of active learning.

## 2 Related Work

### Knowledge Base Population

There is a vast amount of literature devoted to knowledge base population from text, with many established initiatives to foster research on the topic such as the Knowledge Base Population task at TAC^1^ and the TREC Knowledge Base Acceleration track.^2^ In these initiatives, systems are compared on the basis of recognizing individuals belonging to a few selected ontological classes, spanning from the common Person, Place and Organization [37], to more specific classes such as Facility, Weapon, Vehicle [8], Role [28] or Drug [31], among others. Efforts have been made in the biomedical domain to unify the diverse set of ontologies with initiatives such as BioPortal [27] and UMLS [5].

### Biomedical Information Extraction

Extraction of information from biomedical texts is an important area of research for advancing the biomedical sciences. A number of datasets have been proposed which are designed to address specific areas of biomedical natural language processing (BioNLP). One particular organization that has been instrumental in this area is BioCreative; they regularly present challenges to the community in the form of datasets and competitions. The type of challenges they have presented include protein-protein interaction extraction [17, 19], chemical disease relation extraction [41], and chemical protein interaction extraction [18]. The introduction of such datasets is important in that it facilitates the creation of machine learning models for tasks such as entity extraction [22, 10], normalization [42, 22], and relation extraction [39]. The most relevant of these to our work is the chemical protein interaction extraction where the task is to mine text for the existence of relationships between chemicals and proteins i.e. that a particular chemical upregulates or downregulates a protein. However, the creation of such datasets has relied on arduous and expensive labeling by human experts. Our work improves on this by demonstrating that a simple method using active learning is an effective way to quickly create high quality data for the downstream task of knowledge base construction. Our work improves on previous efforts to construct biomedical information extraction datasets by showing that quality data can be obtained quickly on a more limited annotation budget.

### Active Learning

Various approaches have been proposed to minimize the cost of obtaining labelled data, one of the most prominent being distant supervision, which exploits large knowledge bases to automatically label entities in text [3, 12]. Despite being a powerful technique, distant supervision has many drawbacks including poor coverage for tail entities [15], as well as the broad assumption that occurrence of entities can be considered as a positive training example [3]. One way to tackle the problem is to use targeted human annotations to expand the large pool of examples labelled with distant supervision [2] or to perform noise reduction[36].

*Active Learning* aims at acquiring targeted human annotations and using different criteria to identify the best data to annotate next. Some of the most commonly used criteria are: (i) *uncertainty sampling*, which ranks the samples according to the model’s belief it will mislabel them [23]; (ii) *density weighted uncertainty sampling*, which clusters the unlabeled instances to pick examples that the model is uncertain for, but also are “representative” of the underlying distribution [9, 26]; (iii) *QUIRE*, which measures each instance’s informativeness and representativeness by its prediction uncertainty [14]; (iv) Bayesian methods such as *bald* (Bayesian Active Learning by Disagreement) which select examples that maximize the model’s information gain [11] (for a comprehensive survey on active learning we refer the reader to [32]). Finally, in the broad context of natural language processing, deep active learning have proved successful for sentence classification, named entity recognition, and semantic role labeling [34]. Specifically for Named Entity Recognition methods Shen et al. [33] showed that NER models can be trained with active learning and achieve near state of the art performance using significantly less labelled data. Our work builds on classic active learning techniques through a pipelined approach to effectively bootstrap a tool for knowledge base construction while simultaneously creating a high quality dataset for biomedical information extraction.

## 3 BioAct

The foundation of *BioAct* is an active learning pipeline which combines both entity recognition and relation extraction. In the following sections, we detail what this pipeline is and how we applied it to the specific domain of antimicrobial resistance facts.

### 3.1 Active Learning for Knowledge Base Construction

The basic active learning algorithm is shown in Figure 2. At a high level, active learning involves iteratively training a model starting from a small seed dataset which is progressively expanded with new labels for data that are most likely to improve the performance of the model. The seed dataset 𝒟 consists of a set of annotated data which can be acquired using a limited budget, but which is insufficient to effectively train a machine learning model. Along with the small set of labelled data, a large set of unlabelled data 𝒰 is available which is used both to acquire new labels and to probe the model for which samples are most difficult to label. One iteration of active learning then consists of the following:

**Figure 2.**
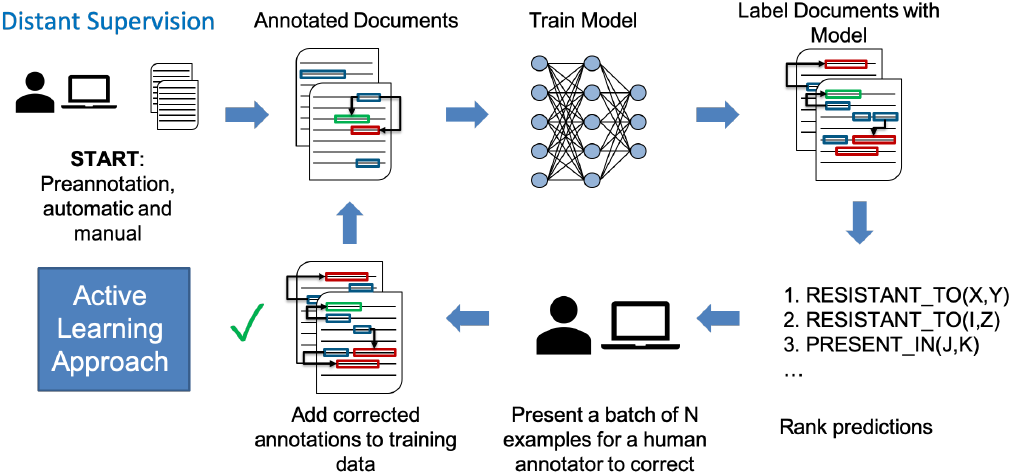
Active learning architecture

- Train the model using the available annotated data 𝒟
- Use the trained model to predict the labels for the data in 𝒰
- Rank the predicted labels in order of their difficulty
- Select the top *N* samples from the ordered predicted labels
- Have a human expert correct the selected predictions
- Add the newly labelled data to the set of labelled data 𝒟

The main considerations are then the source of 𝒰, the source of 𝒟, the model under test, and the method used to rank predictions by their difficulty.

To bootstrap a tool for knowledge base construction, we assume to start with a small manually constructed seed knowledge base 𝒦 consisting of triples (*s, r, o*) ∈ 𝒦. We also assume that each triple in is associated with a set of *j* documents {*ui* |*i* ∈ 1…*j*} ⊂ 𝒰 from which that triple can be extracted. This allows us to build a validation set from the knowledge base 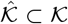 which we can use to gauge the performance of a trained model on the task of knowledge base construction by holding out the documents associated with triples in 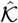 and testing how well a model can recreate 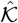 from those documents.

We have a team of experts (13 in this work) label a small subset of 𝒰 to create our labelled seed dataset 𝒟. The tasks we focus on are entity recognition and relation extraction. To this end, we have annotators label the entity types of spans in the documents as well as the relations between pairs of entities. An example labelled sentence is given in Figure 3. The lexicon used is dependent on the problem domain and will be described in subsection 3.2.

**Figure 3.**
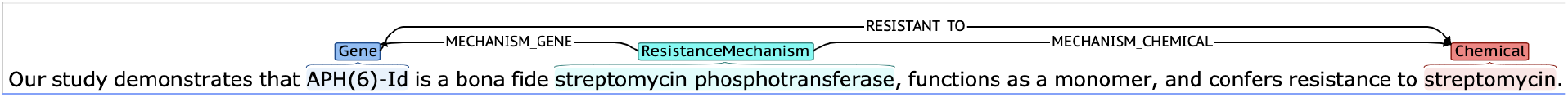
An example labelled sentence.

After obtaining the seed dataset, we reduce the number of human annotators to a small set (3 annotators in this work). We then split the seed dataset into a training set and a small validation set. This data is then used to train a machine learning model on the tasks of entity recognition and relation extraction. The algorithm we use is given in algorithm 1. For relation extraction, the model is trained to predict the class *ŷ*_*rel*_ as the probability of a particular class given the entire input sequence **x**.

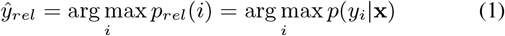

For entity extraction, a linear chain conditional random field (CRF) [20] is used at the output of a deep learning model and the predicted type 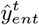 is given as follows.

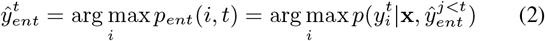

The loss for relation extraction is standard cross entropy loss and for entity recognition we minimize the negative log likelihood of the output of the CRF.

After training the model, we sample a batch of difficult to predict samples for a human annotator to correct. To rank the predictions of our model, we use uncertainty sampling [23]. In this, when we pass a sample through our model, we measure the entropy ℋ of the probability distribution generated by the classifier. For relation extraction, this is simply *Rank*_*R*_ = ℋ (*p*_*rel*_). In the case of entity extraction, we rank sequences by the average of the entropy of the probability distribution of each token in the sequence. In this, for a document of length *T*, the score for entity recognition is given as follows:

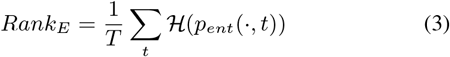

#### Algorithm 1

Active learning algorithm combining entity recognition and relation extraction

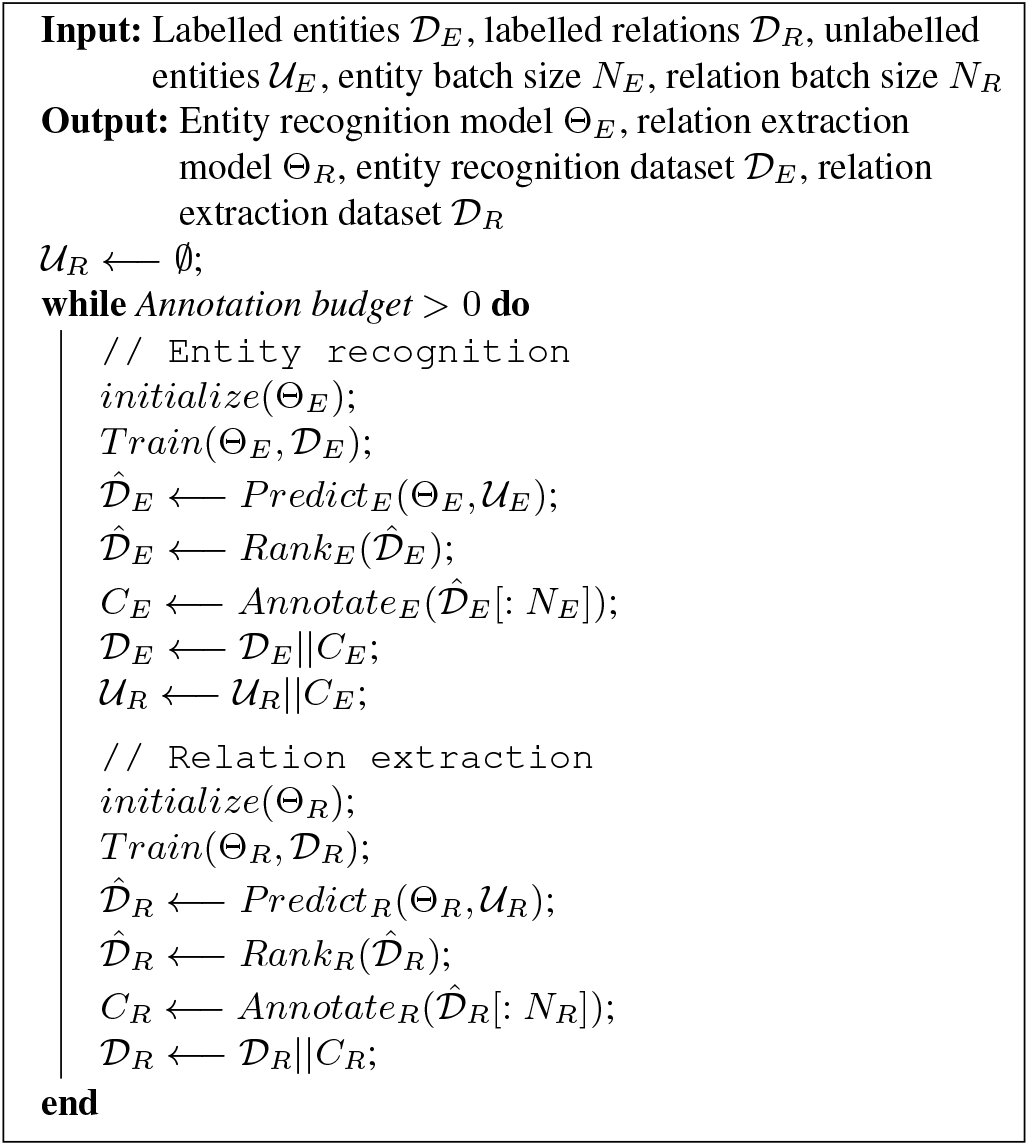

We then use the corrections made by the annotator to measure the F1 score of the model for entity extraction and accuracy for relation extraction. In addition, we test the model on our validation set pulled from the knowledge base to give us insight into how the model improves for the task of knowledge base construction. The newly labelled data is then added to the set 𝒟. This process continues until either model performance reaches a predefined target or the available annotation budget is depleted.

### 3.2 Antimicrobial Resistance Knowledge Base Construction

As a particular use case, we look into the problem of automatically constructing a knowledge base for antimicrobial resistance. This particular domain was selected due to its public health relevance, the existence of manually curated knowledge bases in the domain [16, 21], and the lack of labelled data needed to train and test models for extracting this type of information.

The schema we developed was based on existing work on biomedical information extraction datasets [18, 41, 17, 19] as well as conversations with domain experts in the field. In addition, we chose to incorporate the relations of interest to the community based on existing knowledge bases [16, 21]. The entities and relations we selected are given in Table 1 and Table 2. This schema allows us to capture important high-level antimicrobial resistance facts, being that a particular gene has been shown to confer resistance or be susceptible to a particular chemical. In addition, it allows us to capture which organism the gene was observed in, and the specific mechanism by which the gene confers resistance to a chemical/drug. These latter relation types, as well as the entity types for “ResistanceMechanism” and “ExperimentalTest” have not been previously studied in the biomedical information extraction literature and thus make our dataset novel.

**Table 1.** Entities

| Name | Description |
| --- | --- |
| Gene | A name of a protein/gene/enzyme |
| Chemical | A full chemical name |
| Organism | A full species name, as specific as possible down to the strain level |
| ResistanceMechanism | A pathway or class of enzyme which allows an organism to confer resistance to a particular chemical |
| ExperimentalTest | The name of an *in vitro* experimental test conducted in the study |

**Table 2.** Relations

| Name | Arg1 | Arg2 |
| --- | --- | --- |
| RESISTANT_TO | Gene | Chemical |
| SUSCEPTIBLE_TO | Gene | Chemical |
| PRESENT_IN | Gene | Organism |
| MECHANISM_GENE | ResistanceMechanism | Gene |
| MECHANISM_CHEMICAL | ResistanceMechanism | Chemical |

Full details and descriptions of the entities, as well as our annotation guidelines, are provided in the supplemental material.

### 3.3 Implementation Details

In this work, we considered scientific titles and abstracts as our documents. The initial seed knowledge base was the MegaRes knowledge base [21], which contains triples of the form (Gene, ResistanceMechanism, Chemical). Each triple in the knowledge base has a set of one or more papers associated with it from which the triple can be inferred. In this, our source of unlabelled documents 𝒰 came from MegaRes (as well as the NCBI Pathogen Detection reference gene catalog[6]). We then created the held out validation set 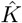 from the MegaRes knowledge base. In addition to MegaRes, we used the CARD knowledge base [16], which is a superset of MegaRes, in our automatic knowledge base construction evaluation (see subsection 4.4).

To obtain annotations from domain experts, we used Brat [35], a common tool for labelling data for information extraction. For the initial labelling phase where we collected the seed dataset, each worker was given a collection of 20 documents and asked to annotate as many as possible within a certain time period (4 weeks). Twice a week we held regular “office hours” to answer any questions workers had regarding annotation, as well as to resolve ambiguous annotations which the workers had trouble resolving. Workers were asked to label both entities and relations during this period. Our annotation guidelines are provided as supplementary material.

During the active learning phase, we reduced the number of workers by approximately 75% (from 13 to 3) and used different tools for collecting entity and relation annotations. For entities, we again used Brat and presented a worker with a batch of full documents to correct on each iteration based on the average entropy of predictions in the document (Equation 3). For relations, we presented the worker with a CSV file where each row contained a single sentence labelled with two entities and a prediction for the relation. The worker was asked to correct the column containing the relation prediction. For entities, we used a batch size of 10 documents and for relations we used a batch size of 100 relations.

For unlabelled relation samples, we were required to have a set of sentences which already had the entities labelled. In this, we corrected the entities in each iteration first, followed by predicting relations on the corrected entities, selecting the top 100 least certain predictions for correction and leaving the remaining (uncorrected samples) in the pool of unlabelled relations.

For our model, we used SciBERT [4], a variant of the BERT [7] model tailored for scientific text. SciBERT is particularly effective for scientific texts as it was pretrained on a large set of scientific papers from Semantic Scholar [1]. We used the default parameters for both the entity recognition variant and text classification variant (for relation extraction). The entity recognition variant of the model uses a Bidirectional LSTM on the output of BERT, as well as a character level CNN. The output is then fed to a linear chain CRF to predict the most likely tag sequence. For relation extraction, we used the text classification model on every pair of entities which co-occur within the same sentence. This essentially boils down to marking the two entities in the sequence with special tokens and predicting the relation type using the [CLS] token which is prepended to every sequence in BERT. The whole pipeline will be made public upon acceptance of the paper.

## 4 Experimental Results

### 4.1 Metrics

We evaluated *BioAct* based on the following criteria

- The amount of labelled data acquired before and after active learning.
- The performance of the trained model on each iteration of active learning. This is evaluated using F1 for named entity recognition and accuracy for relation extraction. Accuracy is used as opposed to F1 since the task given to the subject matter expert is to indicate what the correct relation is for each example. Therefore, accuracy is more appropriate.
- The ability of a model trained using active learning to generalize to unseen relations. We evaluate this using F1 score.
- The ability of a model trained using active learning to incorporate new relations into the knowledge base. We evaluate this using F1 score.

The motivation behind these metrics was to provide insight into the utility of *BioAct* to accomplish the following:

- Build a dataset for biomedical information extraction more quickly than traditional manual curation.
- Train a machine learning model which can augment an existing knowledge base using a dataset constructed through active learning.

### 4.2 Dataset Curation

We first characterize the initial manually curated dataset. Each document was originally annotated by two subject matter experts (SMEs). The initial unlabelled pool of data was collected starting from two sources. The primary source of data was the MegaRes [21] manually curated knowledge base of antimicrobial resistance facts. Each fact in this knowledge base has 0 or more references associated with it. We selected the subset of facts which had at least one reference coming from PubMed or PubMed Central as well as all of the titles and abstracts from these references. This yielded 567 abstracts. The remaining 1007 abstracts were acquired by pulling references associated with entries in the NCBI Pathogen Detection reference gene catalog [6]. We preannotated the genes, chemicals, and organisms in the initial 1,574 documents using PubTator Central [40], which provides an interface for annotating large sets of PubMed and PubMed Central documents using several different text mining and information extraction techniques. From this initial set of documents, 133 of them were doubly annotated within a 4 week period by human experts. The pool of SMEs was recruited through face to face and remote solicitation at the authors’ research institution. An initial set of guidelines was developed, which was later refined after a dry run with a cohort of 5 annotators. The guidelines are included in the supplemental material of this paper. A total of 13 annotators were recruited for the 4 week period. Annotators were given instructions to annotate as many documents as their time allowed during the 4 week period. In retrospect, better incentives were needed in order to improve the number of documents annotated, and it is recommended to build these in beforehand. One critical component of this initial annotation phase was the holding of regular “office hours” where available annotators would meet in a single location. This is where the majority of annotations were made. After the 4 week session, the interannotator agreement (IAA) was calculated using F1 score for both entities and relations. The results for entities are listed in Table 3 and for relations in Table 4.

**Table 3.**
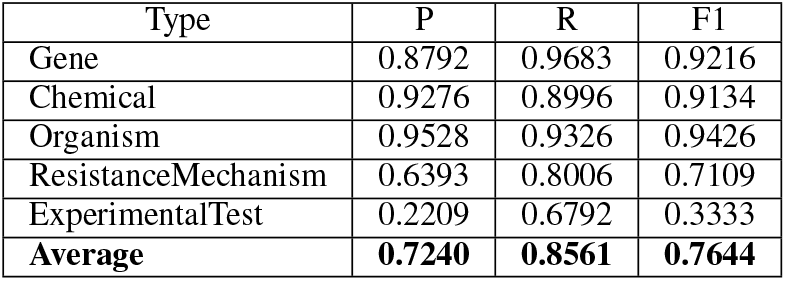
IAA measures for entities

| Type | P | R | F1 |
| --- | --- | --- | --- |
| Gene | 0.8792 | 0.9683 | 0.9216 |
| Chemical | 0.9276 | 0.8996 | 0.9134 |
| Organism | 0.9528 | 0.9326 | 0.9426 |
| ResistanceMechanism | 0.6393 | 0.8006 | 0.7109 |
| ExperimentalTest | 0.2209 | 0.6792 | 0.3333 |
| **Average** | **0.7240** | **0.8561** | **0.7644** |

**Table 4.** IAA measures for relations. Some of the annotators did not annotate any relations, so these documents were omitted from the calculation. There was some confusion on the definitions of the MECHANISM relations at the beginning of annotation, thus driving agreement low. A third expert went through and manually resolved conflicts for every document which was doubly annotated in order to mitigate the annotation issues.

| Type | P | R | F1 |
| --- | --- | --- | --- |
| RESISTANT_TO | 0.4388 | 0.6228 | 0.5149 |
| SUSCEPTIBLE_TO | 0.3571 | 0.3571 | 0.3571 |
| PRESENT_IN | 0.6000 | 0.5091 | 0.5508 |
| MECHANISM_GENE | 0.3889 | 0.1795 | 0.2456 |
| MECHANISM_CHEMICAL | 0.2031 | 0.1831 | 0.1926 |
| **Average** | **0.3976** | **0.3702** | **0.3722** |

We observed relatively high IAA for entities which have been well studied in the literature (genes, chemicals, and organisms). For the new entity type ResistanceMechanism, we saw moderate IAA as there was some initial confusion about how the entity type was defined. ExperimenatalTest saw markedly lower IAA. The reason for this is how annotators decided the length of the phrase that constituted the full experimental test. For example in the sentence “To gain insight into the molecular basis of EMB resistance, we characterized the 10-kb embCAB locus in 16 EMB-resistant and 3 EMB-susceptible genetically distinct Mycobacterium tuberculosis strains from diverse localities by automated DNA sequencing and single-stranded conformation polymorphism analysis,” one annotator labelled “polymorphism analysis” as an experimental test while another labelled “single-stranded conformation polymorphism analysis.”

For relations, we also observed lower IAA. Much of this was due to many annotators only annotating some of the relations in the document. This led to very sparse relation annotations per document. In order to alleviate this, we had a third subject matter expert review each of the 133 documents and resolve conflicts in both entity and relation labels manually.

### 4.3 Active Learning

In all experiments we used SciBERT [4] as the machine learning model, with the entropy of the output classifier used for ranking in uncertainty sampling. We compared the amount of data we collected at the end of active learning with the original manually curated seed dataset. In addition, we compared the amount of time and people it took to collect the data (Table 5). We were able to collect significantly more data in less time and with less human annotators using active learning. This included a 2x increase in the number of entities annotated, and over a 1.5x increase in the number of relations with 25% of the number of annotators.

**Table 5.**
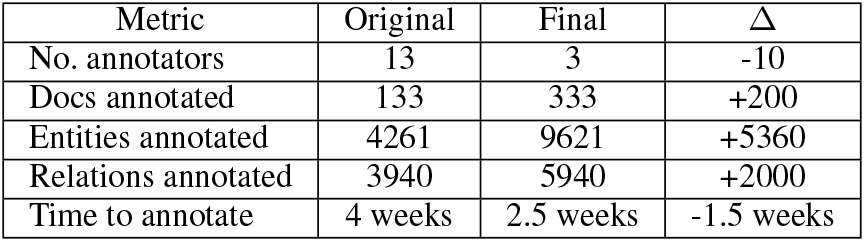
Comparing dataset statistics before and after active learning

Next, we examined how the model performance changed on the data presented to the SME at each iteration of active learning. This tells us how well the model is able to classify its least certain predictions as more data is added to the pool of labelled data. The results for entities (F1) are given in Figure 4 and for relations (accuracy) in Figure 5.

**Figure 4.**
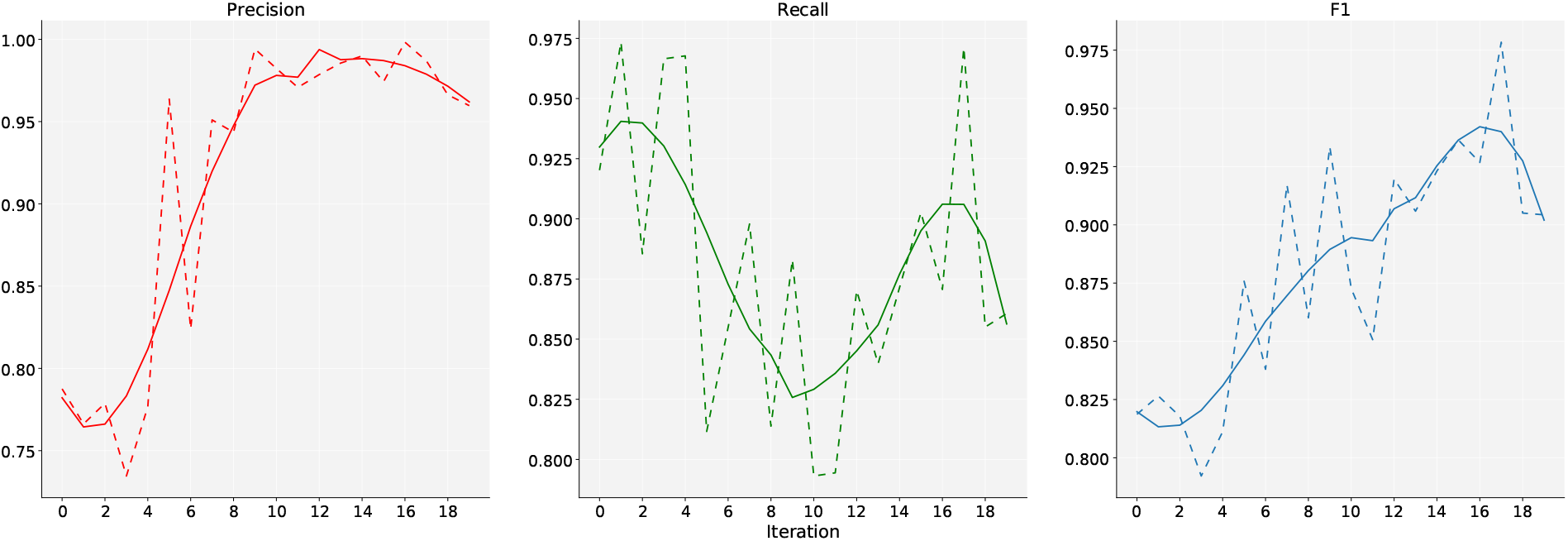
Change in Average F1 score vs. iteration (dotted lines are original data, solid lines are smoothed).

**Figure 5.**
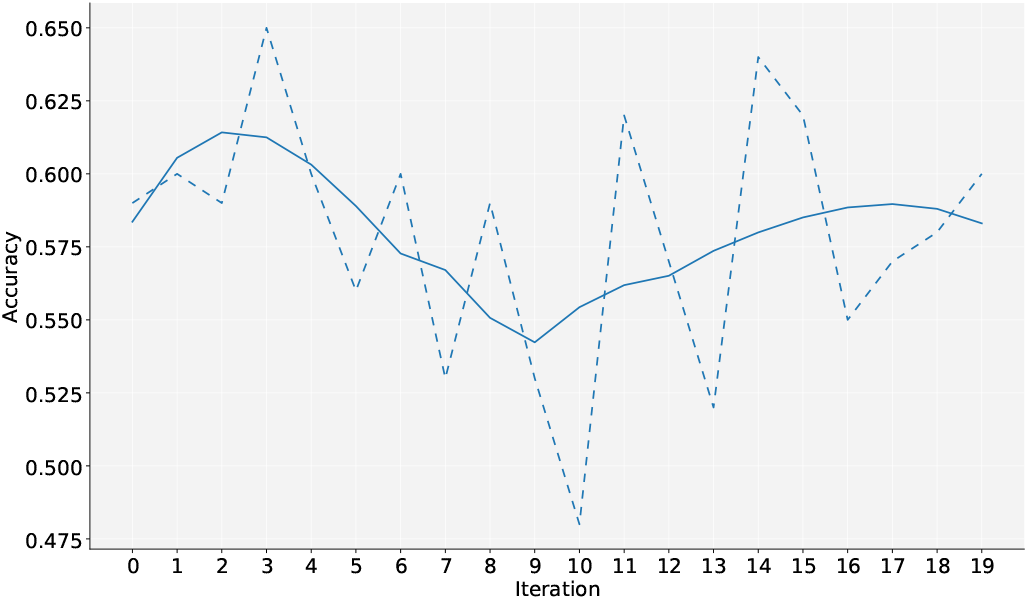
Change in relation extraction accuracy vs. iteration (dotted lines are original data, solid lines are smoothed).

For entities, we saw that the model F1 improved for each iteration of active learning. In particular, the model improved most for the entities which had low IAA in the initial pool of data (Resistance-Mechanism and ExperimentalTest). One potential argument against this is that the model was overfitting to the annotator. Our results on extracting triples will show that, despite this, the model does improve at the primary task of performing automatic knowledge base construction.

Additionally, we saw the biggest gains in terms of the model precision, with recall dropping at first and then rising again. Observing the data which is selected for correction at each iteration of active learning, we saw that the model slowly incorporates new entity types over time. These entities then constitute an increasingly long tail of new entity types that the model is less certain about. As documents are selected by the model which contain these new entities, the recall begins to drop, but the training set is then supplemented with examples of new entities which did not exist in the seed knowledge base (see subsection 4.4). At the same time, the model continues to incorporate examples of entities which are highly represented in the literature since we are selecting whole documents for correction, so the precision progressively increases as the model becomes more readily able to recognize these entities.

For relation extraction, the model initially starts to disagree more with the human annotator, and gradually starts to improve. The reason for this is that the pool of unlabelled relations starts small and grows over time. The set of unlabelled relations at the beginning of learning is unlikely to reflect the distribution of all possible data, so the pool could be easier to predict than average. As more data is added to the pool in each iteration of active learning, more diversity is introduced thus allowing for more potential variation. Once the amount of data becomes large enough, the model begins to improve again since we have incorporated a more diverse set of data on which to train.

### 4.4 Automatic Knowledge Base Reconstruction and Augmentation

The primary purpose of this study was to characterize how a simple active learning setup serves for bootstrapping knowledge base construction in a new domain. In order to accomplish this, we tested the system’s ability to reconstruct and augment a held out portion of a original knowledge base. To acquire this validation data, we randomly selected 200 resistance pairs from the original seed knowledge base (MegaRes [21]) and the set of PubMed articles associated with each pair. From this, we manually inspected each resistance pair and filtered out those which would be impossible to obtain from their associated abstracts. This left us with a validation set of 169 resistance pairs and 228 abstracts. From this, we used the trained model at each iteration of active learning to extract resistance pairs from the 228 validation abstracts. We then used the following metrics to evaluate the performance of the model:

- **Recall** - measured by comparing our extracted relations to the 169 relations which are verified to be attainable from those 228 articles.
- **Precision** - measured by comparing our extracted relations to all of the known relations in both MegaRes and the larger CARD knowledge base [16] from which MegaRes is constructed.
- **F1** - the harmonic mean of the precision and recall calculated above.

As the raw data is not linked to either of those knowledge bases and we do not use a model for entity linking, we manually linked all of the names which our model extracts.

Our results for knowledge base construction are given in Figure 6. This result indicates that *BioAct* can be used to effectively increase the ability of a model to construct a correct knowledge base. The quality of the labels does not degrade, and allows the model to improve its ability to reconstruct the knowledge base. In this, we see relatively consistent precision across iterations with an improvement in recall. This result is encouraging in that it shows that the model mainly incorporates the new names that it is presented as new data is labelled without creating additional false positives.

**Figure 6.**
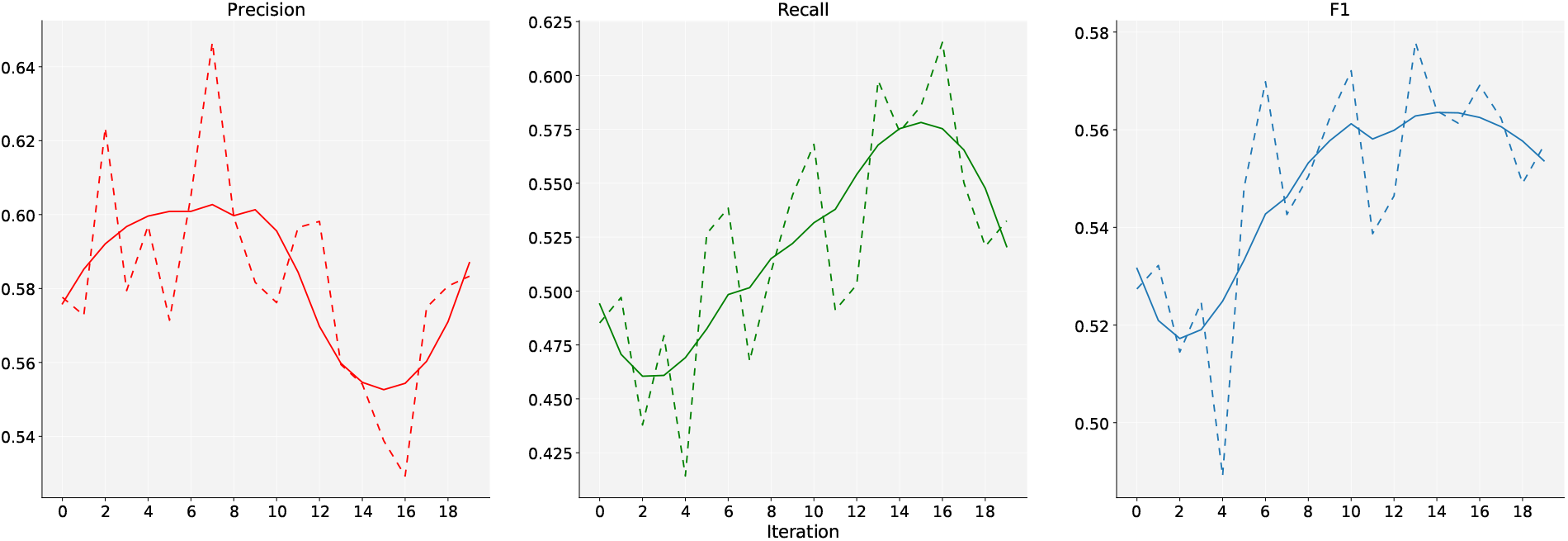
Model F1 performance against source KBs vs. iteration Number of test resistance pairs: 169; Number of test articles: 228

Additionally, we were interested in the number of new relations which the model could add to the source knowledge base. To determine this, we observed the number of relations which were extracted from the test corpus which did not exist in the source knowledge base (MegaRes) but did exist in the larger CARD knowledge base. This gives us an idea of the expected ratio of true positives to false positives in the relations we extract which are unknown to our source knowledge base.

We observed encouraging results in terms of the number of new relations which could be added to the knowledge base. The model continued to find new relations throughout the entirety of training not seen in previous iterations (Figure 7). This result shows that our method can be an effective way to quickly improve a model’s ability to augment an existing knowledge base. However, this should happen in tandem with a human in order to filter out the false positives. Ensuring a higher quality seed corpus could be one way to mitigate this, as our results in Figure 6 indicate that false positives introduced at the start of active learning potentially persist throughout.

**Figure 7.**
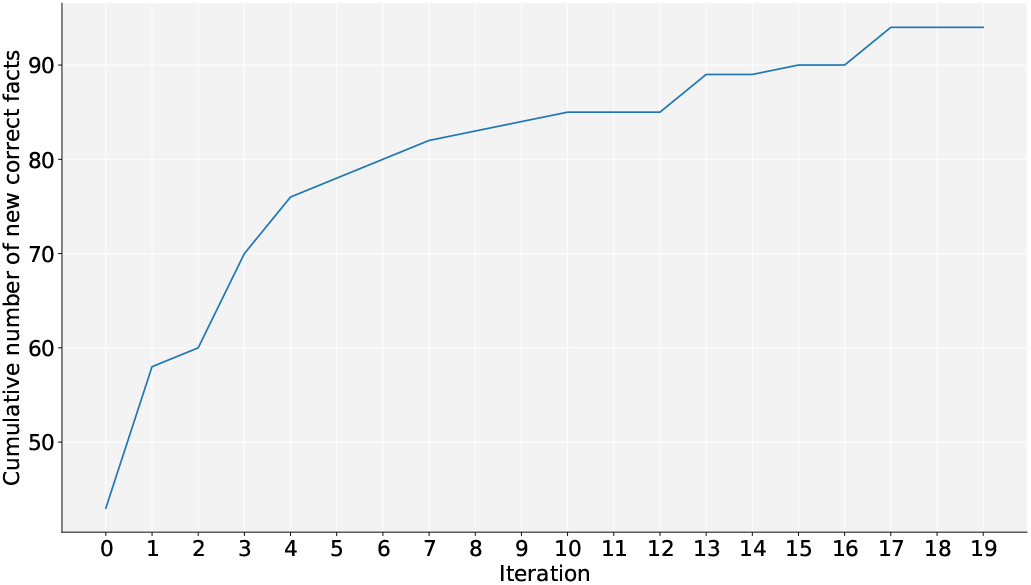
Cumulative new correct relations extracted from the test corpus not in the seed KB vs. iteration

On average we saw the one in three new facts (those that didn’t exist in MegaRes) pulled by our model were actually true positives. This figure is encouraging considering the small amount of data which we had to collect in order to achieve it. Additionally, it indicates that one can effectively collect useful training data with a much smaller cohort of enthusiastic annotators as opposed to relying on the crowd, thus reducing the budget needed for annotation while maintaining annotation quality. Some representative extra facts pulled by the model are given in Table 6, illustrating the type of data a human curator would sift through when correcting new facts. Such a setup would be in line with previous work on never ending learning [24].

**Table 6.** Representative examples of extra resistance relations which the model predicts. **Blue** spans are genes and **red** spans are chemicals.

| Correct | Incorrect |
| --- | --- |
| We show here that the human antimicrobial peptide LL-37 induces the **mefE** promoter and confers resistance to **ery-thromycin** and LL-37. | These **parC** mutations included alterations of serine 80 to arginine or **isoleucine** and glutamate 84 to glycine or lysine. |
| Resistance genes identified in this study were aadA2, aadA4, aadA22, and **aadA23** (for **aminoglycoside** resistance). | The kinetic studies also revealed that **clavulanic acid** and tazobactam inhibited **KPC-1**. |

## 5 Conclusion

In this work we presented *BioAct*, a simple method for quickly building a dataset and tool for biomedical knowledge base construction using active learning. We demonstrated that active learning is useful for rapidly developing a dataset for biomedical information extraction, and that the quality of the data is sufficient to continuously improve the ability of a model to perform knowledge base construction. We focused on the particular domain of antimicrobial resistance and are releasing our data to the community to support research in this important area. While promising, future research is needed into how to mitigate the false positive rate which appears to increase during active learning. In addition, while accelerating the labelling of data in the long run, the method still requires an initial investment in exhaustive manual annotation which could potentially be helped by improved weak supervision using a tool such as Snorkel [29]. We also see the need to incorporate normalization as a component in the active learning pipeline. Finally, there could also be utility in comparing various different methods for ranking predictions, as well as testing different models in order to determine the most effective setup. We hope that this work will inspire further research in this direction, as well as enable research in information extraction of antimicrobial resistance facts.

## Acknowledgements

The authors would like to acknowledge the IBM researchers and domain experts who contributed their time and expertise during the annotation and validation process. This work was completed during an internship at IBM Research Almaden.

## Footnotes

1 http://www.nist.gov/tac/2015/KBP

2 http://trec-kba.org/

