## supplementary materials for "BioAct: Biomedical Knowledge Base Construction using Active Learning"

### Guide for Annotators

#### Why

Resistance to antimicrobial chemicals is becoming an increasingly pervasive problem. 2 million people are affected by antibiotic resistance development leading to 23,000 deaths annually. Efforts to track and catalogue information related to antimicrobial resistance coming from the scientific literature are currently done manually. The ability to automatically create structured data from text about antimicrobial resistance will accelerate these manual curation efforts. To enable this and to encourage research in this area, it will be important to introduce an expert curated dataset of annotated text coming from scientific literature related to antimicrobial resistance.

#### How

The tool that we will be using for creating annotations is **Brat**, an open source web application for adding annotations to text documents. Prior to accessing the server, please briefly look at the Brat manual and keep it on hand while making annotations:
https://brat.nlplab.org/manual.html

**Please make sure to only annotate documents in the collection which you are given (look for your name in the collection browser).**

#### Annotation Guidelines[1]

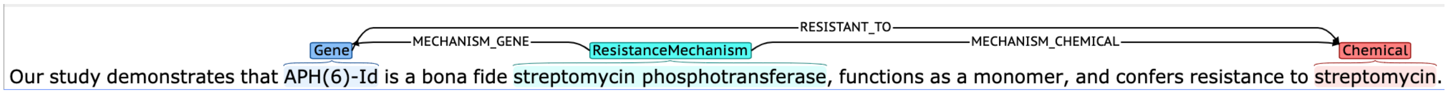


**Figure 1** A sample Gene-Chemical resistance annotation (PMID: 24248535)

This guideline describes the specific types of information that should be annotated in a document to identify facts related to antimicrobial resistance. Annotation of a document consists of labeling selected text with a tag that indicates the type of thing or concept the text represents (e.g. a gene or chemical, as in Figure 1) as well as the relationships that exist between concepts (e.g. that a gene confers resistance to a chemical, also in Figure 1). The term **mention** is used throughout this document to indicate the entities being labeled, e.g., in Figure 1 the text 'streptomycin' is a mention of chemical. The term **relation** is used to indicate the interaction between two mentions (e.g. RESISTANT_TO, PRESENT_IN). For annotation,

[1] These guidelines are largely adapted from
https://tac.nist.gov/2018/SRIE/guidelines/SRIE_AnnotationGuidelines_mentions_05022018.pdf

each type of annotation listed in the sections "Annotating Mentions" and "Annotating Relations" below should be tagged.

### Annotating Mentions

#### Mention Types

**Table 1** Description of Mention types

| Mention Type | Description |
|---|---|
| Gene | A named genetic component. This can be a specific gene **or** protein **or** enzyme name e.g. bla(IMP-4). |
| Chemical | A chemical or drug name e.g. rifampin. |
| ResistanceMechanism | The mechanism that causes a gene to either confer resistance or be susceptible to a chemical e.g. methylation. The mechanism can also be a description of the **gene or protein type** or the **type of enzyme** produced which allows a gene to confer resistance to particular chemicals e.g. streptomycin phosphotransferase |
| ExperimentalTest | Molecular, biochemical, or other experimental test used to confirm results beyond *in silico* methods |
| Organism | The name of an organism e.g. *Escherichia coli* |

#### Mention Annotation Guidelines

- Unless noted in the Guidelines for Specific Mention Types section, annotate the longest contiguous text that describes the item of interest, including abbreviation definitions. Mentions generally should not cross sentence boundaries. Do not include citations at the end of relevant text; citations that appear within the relevant text should be included.

- Only annotate full tokens that can be separated from the rest of the text by punctuation and whitespace

- Annotate only the text parts of a mention, excluding trailing spaces and trailing punctuation (e.g. periods and commas).

- Many mentions will be automatically **pre-annotated** by an external tool. We will need to **correct** any pre-annotated mentions (for example, if the annotation doesn't cover the entire mentioned phrase or the annotated phrase is not actually a mention of the annotated type). Figure 2 illustrates how a sentence may be pre-annotated:

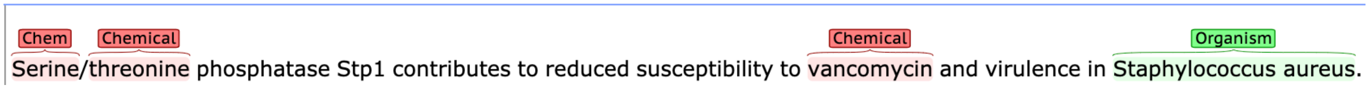


**Figure 2** A pre-annotated sentence which needs to be corrected (PMID: 22492855)

In this example, the mentions "Serine" and "threonine" would need to be deleted and changed to "Serine/threonine phosphatase" and the type would need to be set to "ResistanceMechanism". In addition, the mention "Stp1" would need to be annotated as a Gene, and the relations RESISTANT_TO(Stp1, vancomycin), PRESENT_IN(Stp1, Staphylococcus aureus), MECHANISM_GENE(Serine/threonine phosphatase, Stp1), and MECHANISM_CHEMICAL(Serine/threonine phosphatase, vancomycin) need to be annotated (see Figure 3).

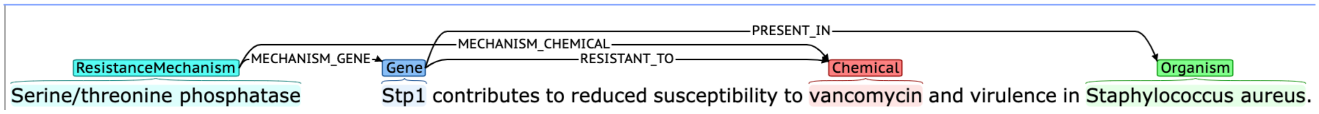


**Figure 3** After annotating the sentence (PMID: 22492855)

To delete a mention, double click on the entity tag. In the pop-up menu, click **Delete**.

To change a the text highlighted for a mention, double click on the entity tag. In the pop-up menu click **Move** and highlight the full text for that mention.

If the annotated mention is not actually of the type indicated, you can either change the type from the pop-up menu or delete the mention entirely if it is not of any of the types in Table 1.

- If an item (often the case for Chemicals and Genes) appears more than once in the text, annotate all instances, including the use of abbreviations.

- Annotate **all** mentions, even those that do not directly apply to the antimicrobial resistance facts conveyed in the document. For instance, if the gene being investigated is 'VanE' still annotate all mentions of 'VanA'. Likewise, label all chemicals even if the document indicates that no resistance is conferred against those chemicals, and all organisms, even if no gene is mentioned which is present in those organisms.

#### Guidelines for Specific Mention Types

- **Gene:** Capture genes, protein, and enzyme names when tagging the "gene" entity type.

- **Chemical:** In addition to named chemicals, annotate all instances of the mention **multidrug** in order to capture multidrug resistance relationships, as well as **antibiotic**.

- **ExperimentalTest:** Capture the full phrase including the noun (what the test was on) and the verb phrase (how the test was conducted) when applicable. Capture the most succinct continuous span of text which describes the test conducted (see Figure 4). Note: experimental tests should be focused on wet lab biological tests opposed to computational-only tests.

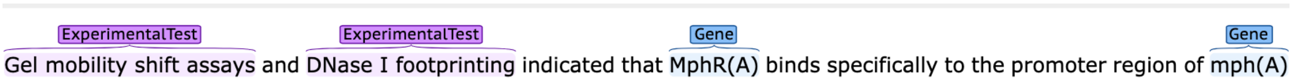


**Figure 4** Two different experimental tests (PMID: 10960087)

- **Organism:** Annotate all formal names (i.e. *Escherichia coli*), abbreviations (i.e. *E. coli*) and common names (i.e. Human) that refer to any organism. Capture the entire name of the organism, including strain if it is given (as in Figure 5). In addition, capture any formal names up to the family level, as well as any text indicating a mutation to an organism. The organism name should be explicit in that it has an actual name (i.e. do not annotate a phrase such as "an unknown species found in…").

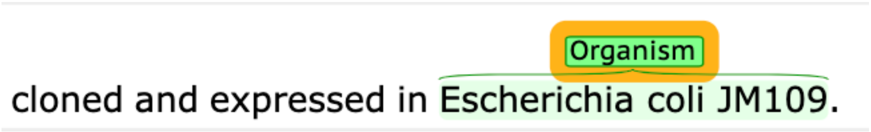


**Figure 5** Strain level annotation (PMID: 10390216)

- **ResistanceMechanism:** Capture all enzymes and pathways listed in the document which explain the means by which a particular gene confers resistance or is susceptible to a particular chemical. This also includes **gene, enzyme, and protein type descriptions** such as "metallo beta-lactamase" which enable a gene to confer resistance to particular chemicals. For example, in Figure 1 we label just "APH(6)-Id" as the gene, and "streptomycin phosphotransferase" as the resistance mechanism since production of this class of enzyme enables resistance against streptomycin.

### Annotating Relations

#### Relation Types

**Table 2** Description of Binary relation types and arguments. 'ARG1' indicates the source mention type (from) and 'ARG2' indicates the target mention type (to). For example, in Figure 1, 'APH(6)-Id' is ARG1 and 'streptomycin' is ARG2. You should first click on ARG1 and drag the mouse to ARG2 when annotating relations.

| Relation Type | ARG1 | ARG2 | Description |
|---|---|---|---|
| RESISTANT_TO | Gene | Chemical | Indicates that the given sentence expresses that a particular gene confers resistance to a particular chemical |
| SUSCEPTIBLE_TO | Gene | Chemical | Indicates that the given sentence expresses that a particular gene is susceptible to a particular chemical |
| PRESENT_IN | Gene | Organism | Indicates that the given sentence expresses that a particular gene is present in a particular organism. |

**Table 3** Description of Ternary relation types and arguments. The relations for resistance mechanisms are **ternary**. If you annotate the MECHANISM_GENE relation you should always annotate the MECHANISM_CHEMICAL relation as well. If both directions cannot be determined from the sentence or adjacent sentences, ignore the relation and just annotate the mention. Examples of resistance mechanism ternary relation are given in Figure 1 and Figure 3.

| Root Mention Type (ARG1) | Relation Type | ARG2 | Description |
|---|---|---|---|
| ResistanceMechanism | MECHANISM_GENE | Gene | The gene which expresses a particular resistance mechanism |

| | | | |
|---|---|---|---|
| | MECHANISM_CHEMICAL | Chemical | The chemical against which a mechanism operates |

#### Relation Annotation Guidelines

- Relationships should be restricted to **within a single sentence or adjacent sentences.** In this, only annotate a relationship if the two mention arguments appear in the same sentence or adjacent sentences and the relationship can be determined using only the text within these sentences.

- Annotate **all** relations for **all** mentions. For example, if a sentence mentions that 5 different genes confer resistance to two different chemicals, create a relation annotation from each gene to each drug (5*2=10 total annotations).

- Only annotate **unambiguous** relations. That is, the sentences must definitively indicate that the relationship between the two mentions exists. For example, for the sentence in Figure 6, we would not annotate the RESISTANT_TO relation between “Mutation A1400G in rss” and “AMK” because the sentence mentions that it was detected in isolates with “diverse MIC [minimum inhibitory concentration] level” to AMK.

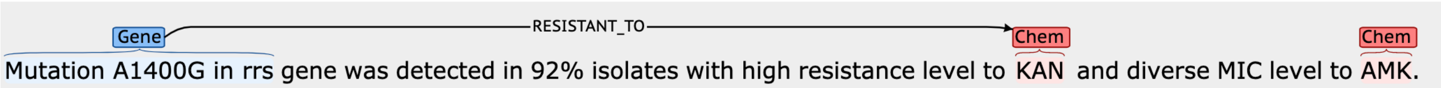


**Figure 6** Example of ambiguous relations (PMID: 25557624)
